# The Effect of Compound Kushen Injection on Cancer Cells: Integrated Identification of Candidate Molecular Mechanisms

**DOI:** 10.1101/503318

**Authors:** Jian Cui, Zhipeng Qu, Yuka Harata-Lee, Hanyuan Shen, Thazin Nwe Aung, Wei Wang, R. Daniel Kortschak, David L Adelson

## Abstract

**Background:** Because Traditional Chinese Medicine (TCM) preparations are often combinations of multiple herbs containing hundreds of compounds, they have been difficult to study. Compound Kushen Injection (CKI) is a complex mixture cancer treatment used in Chinese hospitals for over twenty years.

**Purpose:** To demonstrate that a systematic analysis of molecular changes resulting from complex mixtures of bioactives from TCM can identify a core set of differentially expressed (DE) genes and a reproducible set of candidate pathways.

**Study Design:** We used a cancer cell culture model to measure the effect of CKI on cell cycle phases, apoptosis and correlate those phenotypes with CKI induced changes in gene expression.

**Methods:** We treated cancer cells with CKI in order to generate and analyse high-throughput transcriptome data from two cancer cell lines. We integrated these differential gene expression results with previously reported results.

**Results:** CKI induced cell-cycle arrest and apoptosis and altered the expression of 363 core candidate genes associated with cell cycle, apoptosis, DNA replication/repair and various cancer pathways. Of these, 7 are clinically relevant to cancer diagnosis or therapy and 14 are cell cycle regulators, and most of these 21 candidates are downregulated by CKI. Comparison of our core candidate genes to a database of plant medicinal compounds and their effects on gene expression identified one-to-one, one-to-many and many-to-many regulatory relationships between compounds in CKI and DE genes.

**Conclusions:** By identifying promising candidate pathways and genes associated with CKI based on our transcriptome-based analysis, we have shown this approach is useful for the systematic analysis of molecular changes resulting from complex mixtures of bioactives.

## Introduction

The treatments of choice for cancer are often radiotherapy and/or chemotherapy, and while these can be effective, they can cause quite serious side-effects, including death. These side-effects have driven the search for adjuvant therapies to both mitigate side-effects and/or potentiate the effectiveness of existing therapies. Traditional Chinese Medicine (TCM) is one of the options for adjuvant therapies, particularly in China, but increasingly so in the West. While clinical trial data on the effectiveness of TCM is currently limited, it remains an attractive option because of its long history and because its potential effectiveness is believed to result from the cumulative effects of multiple compounds on multiple targets [13]. Because TCM often has not been subjected to rigorous evidence-based assessment and because it is based on an alternative theoretical system compared to Western medicine, adoption of its plant derived therapeutics has been slow.

In this report, we continue to characterize the molecular effects of Compound Kushen Injection (CKI) on cancer cells. CKI has been approved by the State Food and Drug Administration (SFDA) of China for clinical use since 1995 [31] (State medical license no. Z14021231). CKI is an herbal extract from two TCM plants, Kushen (*Sophora flavescens*) and Baituling (*Smilax Glabra*) and contains more than 200 different chemical compounds. These compounds include alkaloids and flavonoids such as matrine, oxymatrine and kurarinol that have been reported to have anti-cancer activities [31, 47, 35, 48]. Some of these activities have been shown to influence the expression of TP53, BAX, BCL2 and other key genes known to be important in cancer cell growth and survival [45, 38, 25, 17].

We have previously characterized the effect of CKI on the transcriptome of MCF-7 breast carcinoma cells and in this report, we extend our previous results to two additional human cancer cell lines (MDA-MB-231, breast carcinoma and Hep G2, hepatocellular carcinoma). Both cell lines have also been shown to undergo apoptosis in response to the ingredients of CKI [35, 48, 38, 44]. Hep G2 is one of the most sensitive cancer cell lines with respect to exposure to CKI [39] and CKI is often used in conjunction with Western chemotherapy drugs for the treatment of liver cancer patients in China. While the specific mechanism of action of CKI is unknown, several recent studies have reported that CKI or its primary compounds affect the regulation/expression of oncogene products including *β*-catenin, TP53, STAT3 and AKT [31, 35, 22, 18, 41].

However, these and other reports did not evaluate the entire range of molecular changes from treatment with a multi-component mixture such as CKI [10, 8]. Whilst several research databases and tools for TCM research have been developed [32, 34, 5], they are limited by the fact that most of the studies that contribute to the corpus of these databases are from different experimental systems, use single compounds or measure effects based on one or a handful of genes/gene products.

In contrast to previous studies, our strategy was to carry out comprehensive transcriptome profiling and network reconstruction from cancer cells treated with CKI. Instead of focusing on specific genes or pathways in order to design experiments, we have linked phenotypic assessment and RNA-seq analysis to CKI treatment. This allows us to present an unbiased, comprehensive analysis of CKI specific responses of biological networks associated with cancer. Our results indicate that different cancer cell lines that undergo apoptosis in response to CKI treatment can exhibit different CKI induced gene expression profiles that nonetheless implicate similar core genes and pathways in multiple cell lines.

The current study presents the effects of CKI on gene expression in cancer cells with an aim to identify candidate pathways and regulatory networks that may be perturbed by CKI *in vivo*. To this end we primarily use concentrations of CKI higher than used *in vivo* in order to be able to detect effects in the short time frames available to tissue culture experiments. We also combine our current analysis with previously published data to focus on a shared, much smaller set of candidate genes and pathways.

## Material and Methods

### Cell culture and reagents

CKI (total alkaloids concentration of 25 mg/ml) in 5 ml ampoules was provided by Zhendong Pharmaceutical Co. Ltd. (Beijing, China). Chemotherapeutic agent, Fluorouracil (5-FU) was purchased from Sigma-Aldrich (MO, USA). A human breast adenocarcinoma cell line, MDA-MB-231 and a hepatocellular carcinoma cell line Hep G2 were purchased from American Type Culture Collection (ATCC, VA, USA). The cells were cultured in Dulbecco’s Modified Eagle Medium (Thermo Fisher Scientific, MA, USA) supplemented with 10% fetal bovine serum (Thermo Fisher Scientific). Both cell lines were cultured at 37°C with 5% CO_2_.

For all *in vitro* assays, 4×10^5^ cells were seeded in 6-well trays and cultured overnight before being treated with either CKI (at 1 mg/ml and 2 mg/ml of total alkaloids) or 5-FU (150 µg/ml for Hep G2 and 20 µg/ml for MDA-MB-231). As a negative control, cells were treated with medium only and labelled as “untreated”. After 24 and 48 hours of treatment, cells were harvested and subjected to the downstream experiments.

### Cell cycle and apoptosis assay

The assay was performed as previously described [28]. For each cell line, three operators replicated the assay twice in order to ensure reproducibility of the observations. The results were obtained by flow cytometry using either FACScanto or LSRII (BD Biosciences, NJ, US).

### RNA isolation and sequencing

The treated cells were harvested, and the cell pellets were snap frozen with liquid nitrogen and stored at −80°C. Total RNA was isolated with PureLink™ RNA Mini Kit (Thermo Fisher Scientific) according to the manufacturer’s protocol. After quantified using a NanoDrop Spectrophotometer ND-1000 (Thermo Fisher Scientific), the quality of the total RNA was verified on a Bioanalyzer by Cancer Genome Facility (SA, Australia) ensuring all samples had RINs>7.0.

For both cell lines, the sequencing was performed in Ramaciotti Centre for Genomics (NSW, Australia). The sample preparation for each cell line was TruSeq Stranded mRNA-seq with dual indexed, on the NextSeq500 v2 platform. The parameter was 75bp paired-end High Output. The fastq files were generated and trimmed through Basespace with application FASTQ Generation *v1.0.0*.

### Bioinformatics analysis of RNA sequencing

The clean Hep G2 reads were aligned to reference genome (hg38) using STAR v2.5.1 with following parameters: –outFilterMultimapNmax 20 –outFilterMismatchNmax 10 –outSAMtype BAM SortedByCoordinate –outSAMstrandField intronMotif [6]. The clean MDA-MB-231 reads were aligned to reference genome (hg19) using TopHat2 v2.1.1 with following parameters: –read-gap-length 2 –read-edit-dist 2 [15]. Differential expression analysis for reference genes was performed with edgeR and differentially expressed (DE) genes were selected with a False Discovery Rate<0.05 [29].

The DE genes in common for both Hep G2 and MDA-MB-231 cell lines at 24 hours and 48 hours after CKI treatment were selected as “shared” genes. These shared genes were utilized to describe the major anti-cancer functions and principal mechanisms of CKI.

Gene Ontology (GO), and Kyoto Encyclopedia of Gene and Genomes (KEGG) over-representation analyses of both cell lines were carried out using the online database system ConsensusPathDB [14] with the following settings: “Biological process” at third level (for GO); q values (<0.01) were corrected for multiple testing with the system default settings. Disease ontology (DO) over-representation analyses of both cell lines were performed by using the Bioconductor R package clusterProfiler v3.5.1 [40]. For the function analyses of shared/core genes, the method was as similar as our previous study [28] using ClueGO app 2.2.5 in Cytoscape v3.6.0. We enriched our GO terms in the biological process category level 3 and KEGG pathways, showing only terms/pathways with *p* values less than 0.01. Specific Over-represented terms/pathways and gene expression status mapping in KEGG pathways were visualised with the R package “Pathview” [21].

### Gene expression-based investigation of bioactive components in CKI

To integrate with previous data from MCF-7 cells [28], all the shared DE genes regulated by CKI identified in all three cell lines using edgeR were mapped to the BATMAN-TCM database [19]. The pharmacophore modelling method [16] was used to generate the interaction network between the key genes and TCM components using R package igraph [4].

### Reverse transcription quantitative polymerase chain reaction (RT-qPCR)

RT-qPCR was performed as previously described 28]. The list of target genes selected for this study and the sequences of all primers are shown in Additional file 1: Table S1.

## Results

### Effect of CKI on the cell cycle and apoptosis

In our previous study, CKI significantly perturbed/suppressed cancer cell target genes/networks. In the current study we present results that confirm and generalize our previous work. We observed in the MCF-7 study, low concentrations of CKI in our short-term cell assay showed no/little phenotypic effect within 48 hours, and very high doses resulted in excessive cell death at 48 hours precluding the isolation of sufficient RNA for transcriptome analysis [28]. Therefore, in our current study with the two additional cell lines, to ensure consistency, we also selected 1 mg/ml and 2 mg/ml total alkaloid concentrations of CKI for our assays because they generated reproducible and significant phenotypic effects in our cell culture assay.

We used flow cytometric analysis of propidium iodide stained cells to assess both CKI induced alterations to the cell cycle and apoptosis. In Hep G2 cells, CKI treatment resulted in an overall increase in the proportion of cells in G1 phase and decrease in S phase (Fig. 1a and b). Similarly, in MDA-MB-231 cells, although a consistent increase in G1 phase was not observed, CKI caused a decrease in S phase particularly at the 24-hour time point (Fig. 1a and b) indicating possible incidence of cell cycle arrest at G1 phase. Furthermore, at 2 mg/ml of total alkaloids, CKI consistently induced significantly higher level of apoptosis in both cell lines at both time points compared to untreated controls (Fig. 1c). These data together suggest that CKI has effects on the cell cycle by interfering with the transition between G1 to S phase as well as by acting on the apoptosis pathway and promoting cell death.

**Figure 1.**
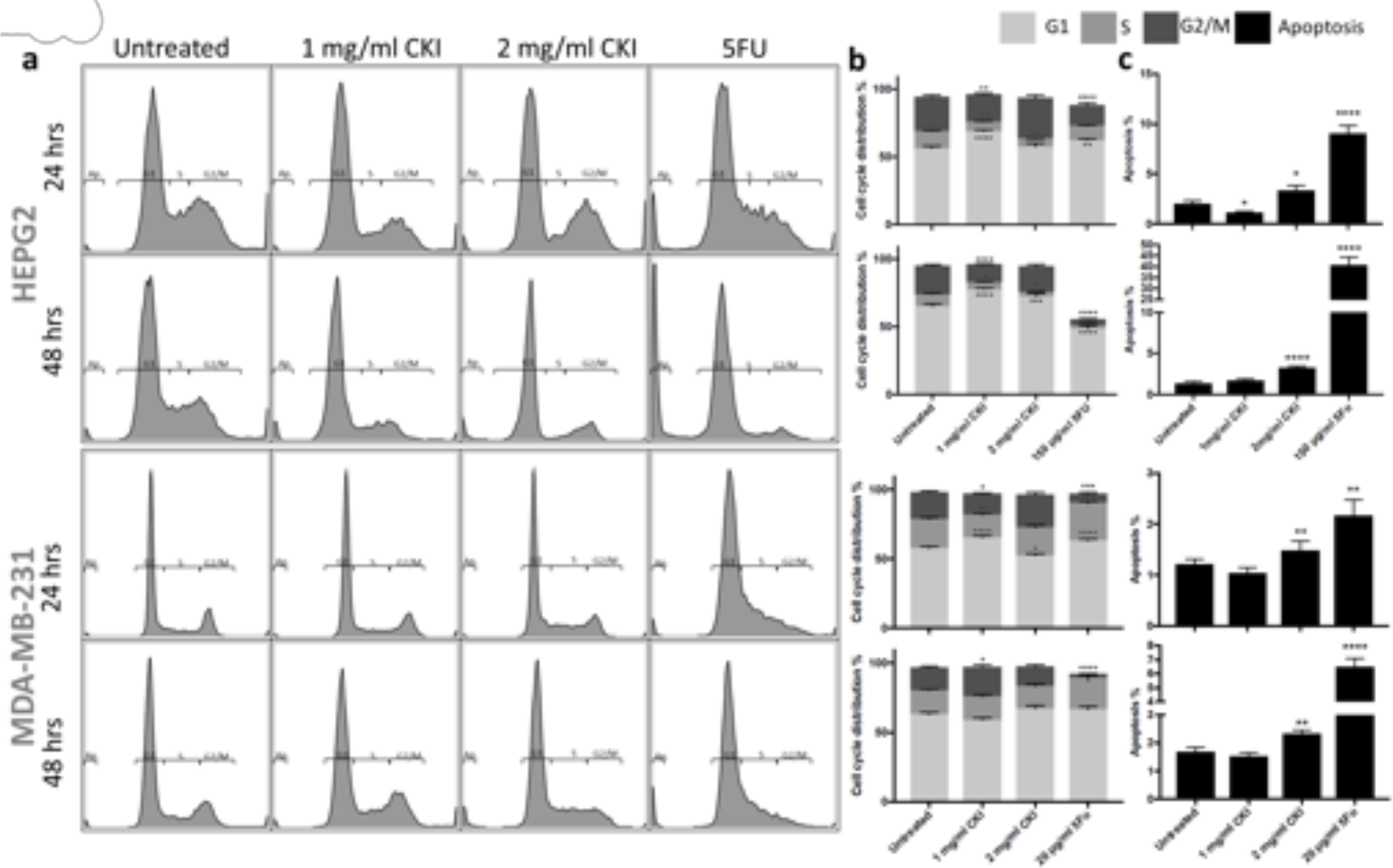
Effects of different treatment on cell cycle and apoptosis of Hep G2 and MDA-MB-231 cells. A) The apoptosis and cell cycle distribution of each cell line after 24- and 48-hour treatments with CKI or 5-Fu assessed PI staining. B) Percentages of cells in different phases of cell cycle resulting from treatment. c Percentage of apoptotic cells after treatment. Results shown are mean ±SEM (n=6). Statistically significant differences from untreated control were identified using two-way ANOVA (*p<0.05, **p<0.01, ****p<0.0001).

### CKI perturbation of gene expression

In order to elucidate the molecular mechanisms of action of CKI on these cancer cells, transcriptome analysis of CKI treated cells was performed. As mentioned above, RNA samples from two cell lines were sequenced with 2×75 bp paired-end reads. We had previously sequenced transcriptomes from CKI treated MCF-7 cells [28] and have included those results for comparison below. The samples from each cell line contained 7 groups at 3 time points (Fig. 2a), in triplicate for every group. In the multidimensional scaling (MDS) analysis, each cell line clustered independently and generally, within the cell line clusters, untreated cells clustered apart from treated cells (Additional file 2: Fig. S1).

**Figure 2.**
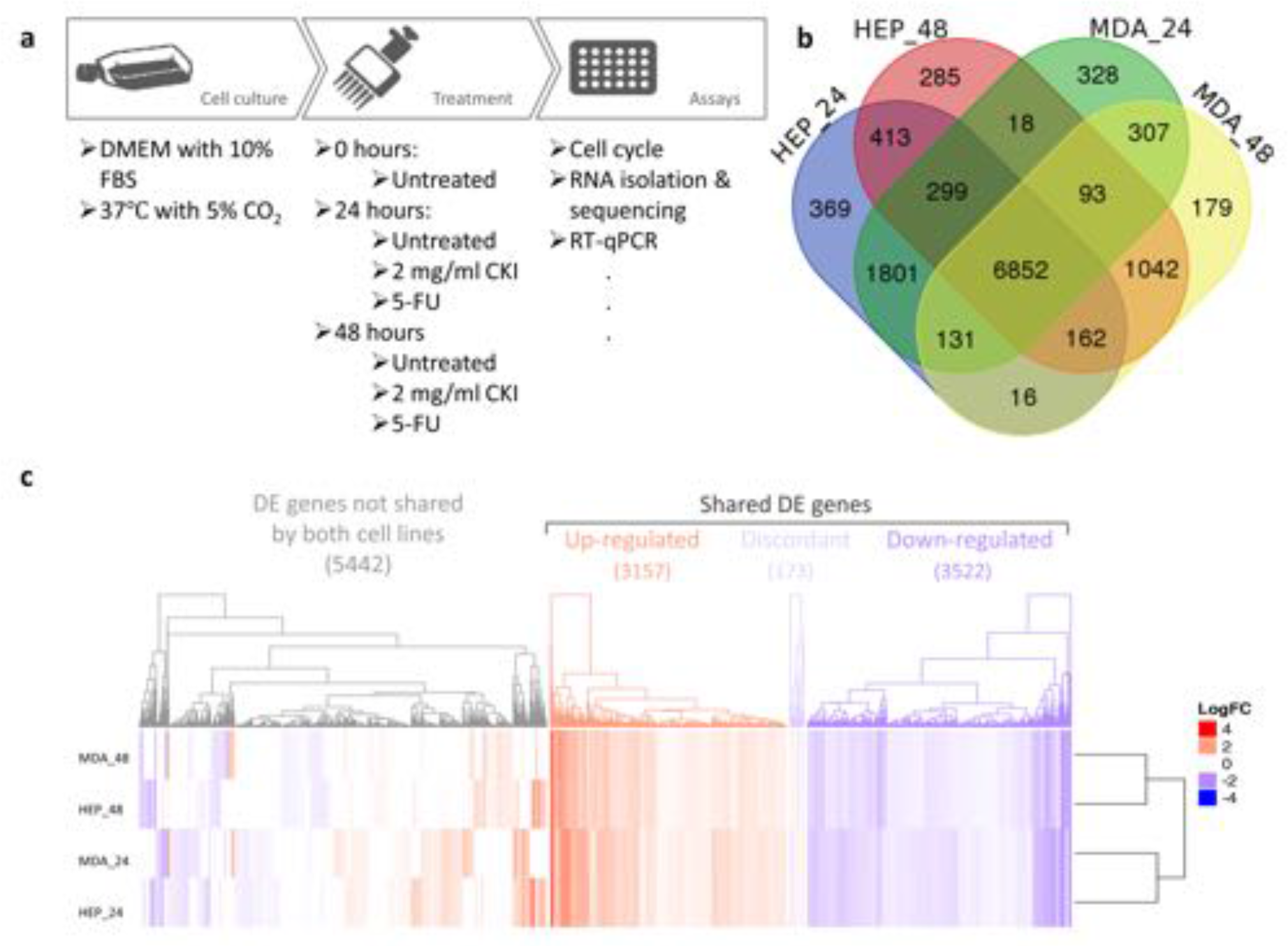
DE genes shared in both cell lines at both time points. A) Work flow diagram showing experimental design and sample collection. B) Venn diagram showing the number of shared DE genes between Hep G2 and MDA-MB-231. C) Heatmap presenting the overall gene expression pattern in both cell lines treated with CKI. Heatmap is split into four parts based on gene content and expression pattern: 5442 differentially regulated genes with expression not shared between the two cell lines, 3157 upregulated genes shared between both cell lines, 3522 down-regulated genes shared between both cell lines, and 173 discordantly regulated genes with differential expression shared between both cell lines.

With the mapping rate were around 90% (Additional file 3: Table S2), a p-value based ranked list of DE genes (compared to untreated from each time point) was generated for both cell lines (Additional file 4: Table S3, sheet 1-4). This list was used to select the shared DE genes. This analysis generated thousands of DE genes (Additional file 4: Table S3, sheet 5) across Hep G2 and MDA-MB-231 cell lines.

Because for each cell line the respective treatment groups clustered together on the MDS plot, there were large numbers of shared genes between them. As a result, we identified a set of 6852 shared DE genes by identifying common DE genes from Hep G2 and MDA-MB-231 cell lines, at 24 hours and 48 hours (Fig. 2b). These shared genes might predict a common molecular signature for CKI’s activity. However, there were still a large number of DE genes that were not shared by both cell lines, as seen in the heatmap in Fig. 2c. The expression of the shared gene set in both Hep G2 and MDA-MB-231 is highly consistent. Interestingly, this consistency is with respect to treatment time, rather than with respect to cell line.

### RT-qPCR validation and dose response of gene expression to CKI

Based on our previous results [28], and analysis below, we selected the 4 top ranked DE genes expressed in G1-S phase of the cell cycle (TP53 and CCND1 for expression level validation and E2F2 and PCNA for low dose response), as well as the proliferation and differentiation relevant ras subfamily encoding gene (RAP1GAP1) for low dose response. We also selected a prominently expressed gene (CYP1A1) for validation because of its sensitivity to CKI treatment. CYP1A1, TP53 and CCND1 expression changes were validated with RT-qPCR with all three genes showing similar patterns of expression in the transcriptome data and RT-qPCR (Fig. 3a).

**Figure 3.**
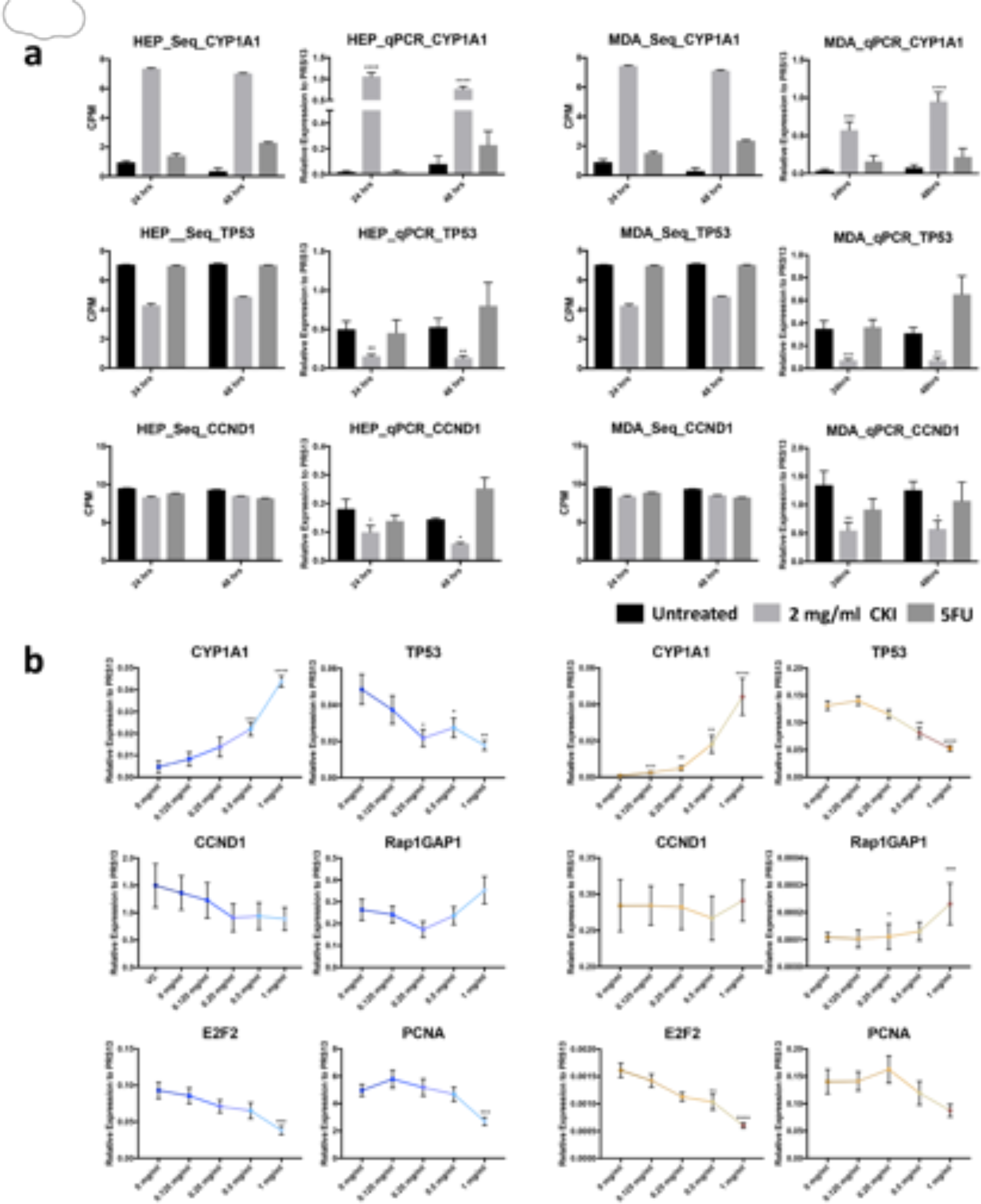
Validation of gene expression and effects of low dose CKI using RT-qPCR. A) Comparison of DE genes between RNA-seq results (left) and RT-qPCR validation (right) for each cell line at 2 time-points. Three DE genes (CYP1A1, TP53 and CCND1) were chosen for validation. Gene expression was generally consistent between transcriptome data and qPCR data. B) Dose response of CKI using a subset of genes with conserved expression in Hep G2 (left), and MDA-MB-231 (right) from 0 mg/ml to 1 mg/ml of total alkaloids. Six genes (CYP1A1, TP53, CCND1, Rap2GAP1, E2F2 and PCNA) were selected based on their relevance to important pathways perturbed by CKI. RT-qPCR results are presented as expression relative to RPS13. Data are represented as mean ±SEM (n>3). A t-test was used to compare CKI doses with “untreated” (*p<0.05, **p<0.01, ***p<0.001).

Because low dose treatment with CKI did not cause significant gross phenotypic effects in either cell line, we decided to use gene expression as a more sensitive measure of phenotype to look at the effect of lower doses of CKI. We used 0.125 mg/ml, 0.25 mg/ml, 0.5 mg/ml and 1 mg/ml concentrations to look for dose dependency of gene expression. Our results showed an obvious dose-dependent expression trend (Fig. 3b) in both cell lines. Because the 0.125 mg/ml concentration of CKI is equivalent to what cancer patients are treated with, our results are potentially clinically relevant.

### Function enrichment analysis

To identify candidate mechanisms of action of CKI, we carried out functional enrichment analysis. We used ConsensusPathDB [14] and Clusterprofiler [16] along with GO and KEGG pathways for over-representation analysis, along with disease ontology (DO) [30] enrichment.

GO over-representation test was determined based on Biological Process level 3 and q value <0.01. The results for both cell lines at both time points were summarized and visualized based on semantic analysis of terms in Fig. 4a. From this result, it was obvious that there were a large proportion of enriched GO terms relating to cell cycle, such as “cell cycle checkpoint”, “negative/positive regulation of cell cycle process” and so on prominently featured for all data sets (Additional file 5: Fig. S2, Additional file 6: Table S4, sheet 1-4).

**Figure 4.**
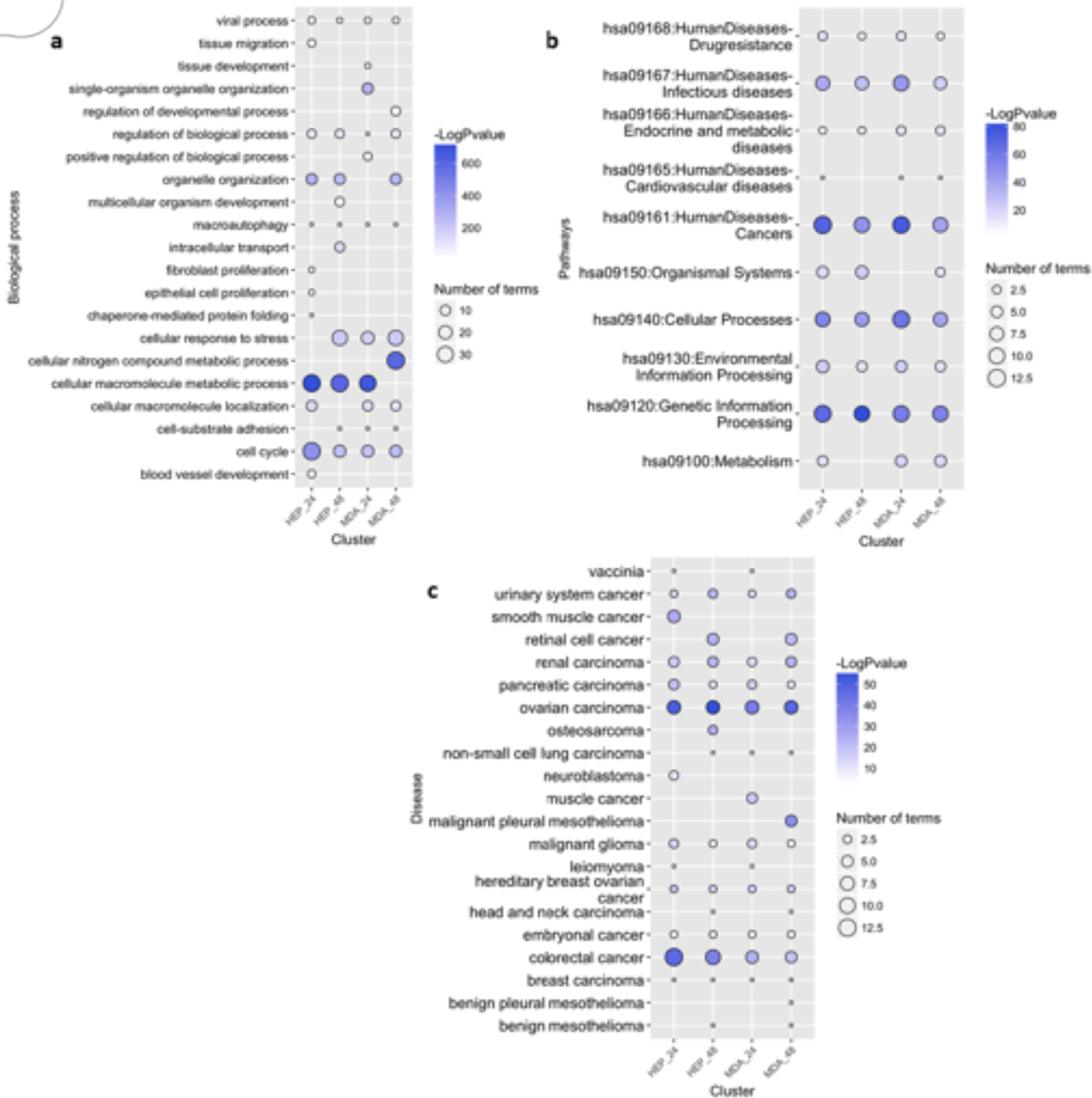
Functional annotation of DE genes for each cell line as a result of CKI treatment. Summary of over-represented A) GO terms for Biological Process, B) KEGG pathways and C) DO terms for DE genes as a result of CKI treatment in each cell line at two time points. For GO semantic and enrichment analysis, Lin’s algorithm was applied to cluster and summarize similar functions based on GO terms found in every treatment. Similarly, by back-tracing the upstream categories in the KEGG Ontology, we were able to obtain a more generalized summary of KEGG pathways for each treatment. The size of each bubble represents the number of GO terms/pathways, and the colour shows the statistical significance of the relevant function or pathways. The DO summary for each treatment was determined by back-tracing to parent terms.

We then used KEGG pathways to determine the specific pathways altered by CKI in cancer. The most regulated over-representative KEGG pathways are summarized according to KEGG Orthology (KO) (Fig. 4b). Cell cycle related pathways such as “cell cycle”, “DNA replication”, and “apoptosis” were also consistently seen in the KEGG enrichment results (Additional file 6: Table S4, sheet5-8) at both 24 and 48 hours. Moreover, in addition to the cell cycle relevant pathways, some cancer related pathways were also observed, such as “prostate cancer” and “chronic myeloid leukaemia”, and a large number of DE genes (283) from the two cell lines were relevant in “pathways in cancer”.

Because the KEGG enrichment revealed many pathways relating to diseases, most of which were cancers, we decided to explore the enrichment of DE genes with respect to DO terms (Fig. 4c). In the DO list (Additional file 6: Table S4, sheet 9-12), all top ranked terms listed are cancers. Interestingly, most cancer types listed are from the lower abdomen, for example “ovarian cancer”, “urinary bladder cancer “and “prostate cancer” etc. occurring in genitourinary organs (Additional file 6: Table S4, sheet 9-12). For both KEGG pathway and DO enrichment, the effects of CKI on both cell lines were similar.

In addition to cell line specific functional enrichment of DE genes, we also analyzed the over-represented GO terms for shared DE genes (Fig. 5a). The most significant clusters were highly relevant to metabolic process, such as “cellular macromolecule metabolic process”, as well as the corresponding positive/negative regulatory biological process (Additional file 6: Table S4, sheet 13). Moreover, various signaling pathways, though not forming a large cluster, were also significant, for example, “regulation of signal transduction” and “intracellular receptor signaling pathway”. Finally, some “cell cycle” related terms constituted relatively large sub-clusters, including “cell division” and “mitotic cell cycle process”. The enriched GO analysis was consistent with the cell line specific enriched results, and with our previous analysis of MCF-7 cells [28]. It is worth noting that for “cell cycle” related terms, most of the participating genes were down-regulated by CKI.

**Figure 5.**
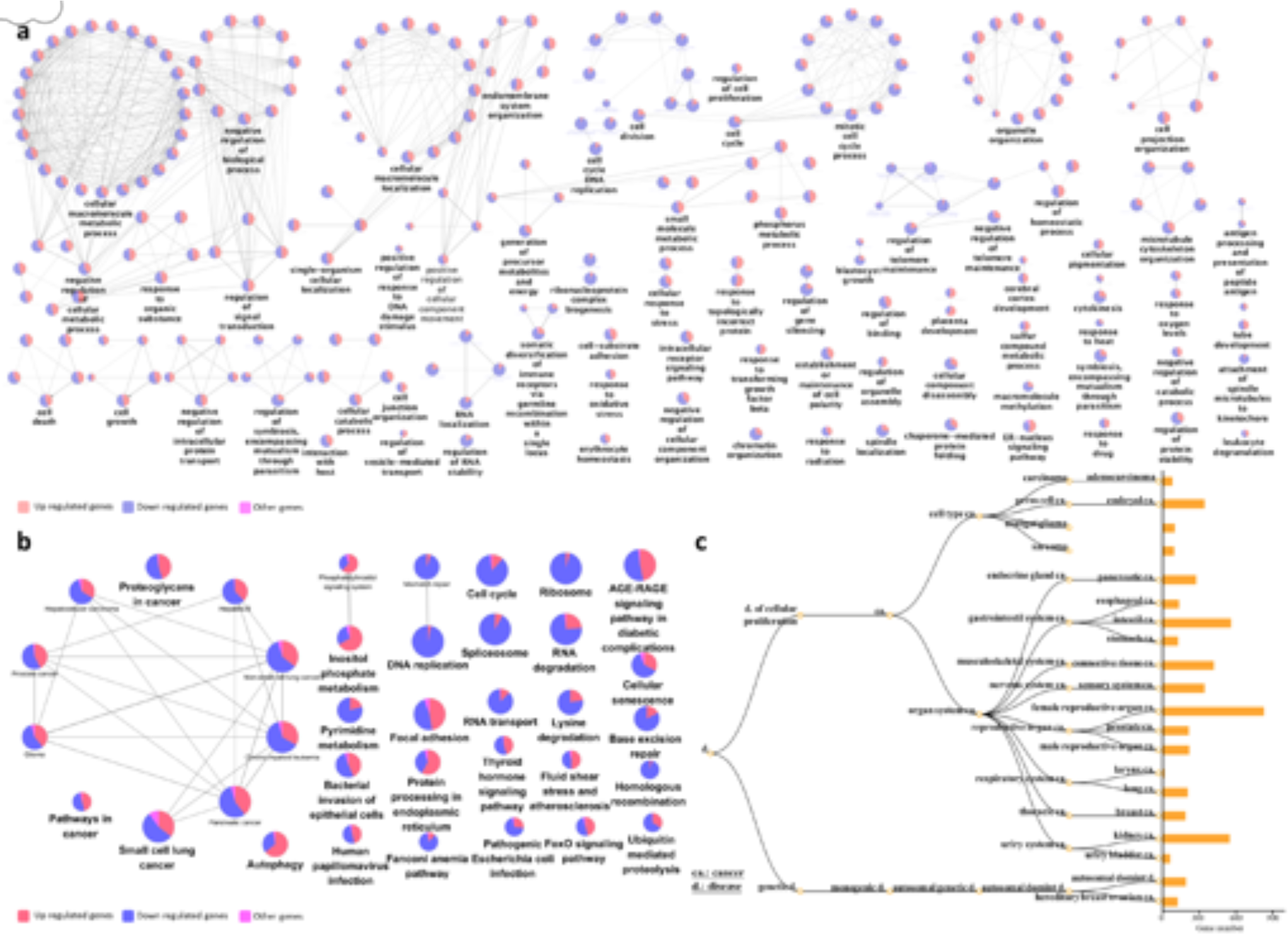
Functional annotation of DE genes with shared expression in both cell lines as a result of CKI treatment. Over-representation analysis was performed to determine A) GO terms for Biological Process, B) KEGG pathways, and C) DO terms for DE genes shared in both cell lines. In nodes for both GO terms and KEGG pathways, node size is proportional to the statistical significance of over-representation. For DO terms, all the enriched terms are statistically significant (p<1×10^-5^) in each category, and the bar length represents the number of expressed genes that map to the term.

Similar results were observed from KEGG analysis (Fig. 5b, and Additional file 6: Table S4, sheet 14) of shared genes. Various pathways related to cancer, formed a large cluster. Pathways such as “DNA replication”, “Ribosome” and “cell cycle” were mostly down-regulated, while up-regulated pathways included “inositol phosphate metabolism” and “protein processing in endoplasmic reticulum”.

We also carried out over-representation analysis of DO terms (Fig. 5c) for all shared DE genes. The analysis results were consistent with the single cell line DO term analysis with mostly cancer related terms; in particular genitourinary or breast cancer terms. While this was also partially similar to the KEGG results for shared DE genes, there were some differences in the KEGG results for disease pathways compared to the DO results, such as “bacterial invasion of epithelial cells”, “Fanconi anemia pathway” and “AGE-RAGE pathway in diabetic complications”.

Specific to the therapeutic potential of CKI for cancer treatment, we applied our data set mapping to KEGG cancer pathways: pathways in cancer - homo sapiens (Additional file 7: Fig. S3). The R package Pathview [21] was used to integrate log fold change values of all the gene expression patterns into these target pathways. Within the 21 pathways in cancer, the “cell cycle” still featured prominently (Fig. 6a). The expression of almost every gene in the cell cycle pathway was affected by CKI, with most of them suppressed. We did not observe this kind of overall pathway suppression in any of the other pathways. We have displayed the summaries for the remaining 20 pathways in the heatmap in Fig. 6b. Although all the pathways were all perturbed by CKI, they include both over and under expressed genes in roughly equal proportions.

**Figure 6.**
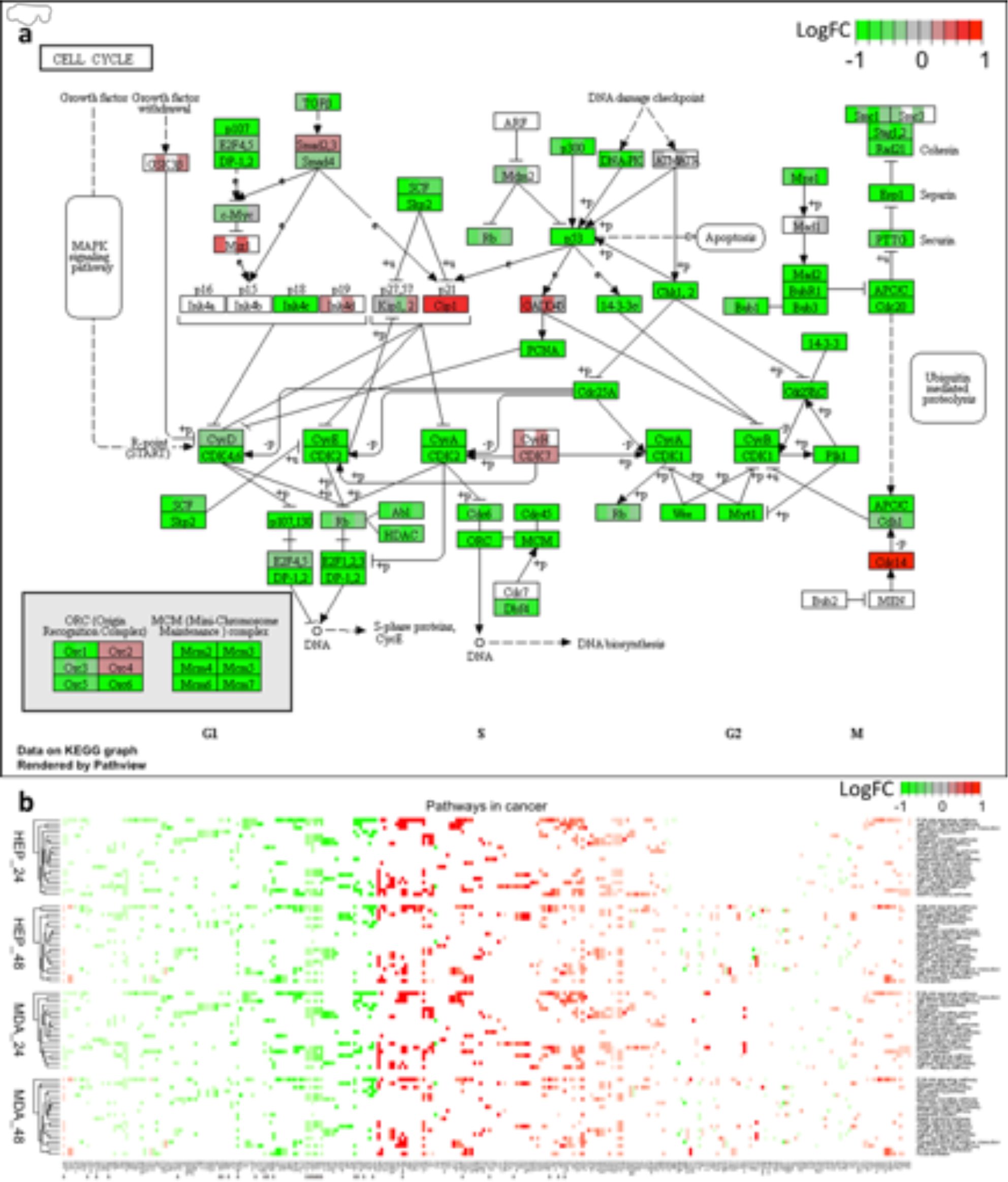
Comparison of shared genes expression in specific pathways across two cell lines. A) Cell cycle pathways, where each coloured box is separated into 4 parts, from left to right representing 24 hour CKI treated Hep G2, 48 hour CKI treated Hep G2, 24 hour CKI treated MDA-MB-231 as well as 48 hour CKI treated MDA-MB-231. B) Heatmap of pathways in cancer. The top two heatmaps summarise the effects of CKI on Hep G2 cells for two time-points, and the bottom two heatmaps show the effects of CKI on MDA-MB-231 cells. In addition to the cell cycle pathway, there were 21 associated pathways in cancer that were perturbed by CKI. The effects of CKI on both cell lines were similar, with changes in TARGET database genes indicated by arrows. Compared to other pathways in cancer, the effects of CKI on the cell cycle pathway showed overall down-regulation.

Collectively, these results suggest a direct anti-cancer effect of CKI, and implicate specific candidate mechanisms of action based on the perturbed molecular networks. The most obvious example is the cell cycle, where G1-S phase is significantly altered, resulting in the induction of apoptosis. The downstream process triggered by CKI is the suppression of gene expression of cell cycle regulators, including TP53 and CCND1. The other perturbed cancer pathways provide additional candidate mechanisms of action for CKI. In the following section we integrate these results with previous results reported in the literature to refine the core set of genes and pathways perturbed by CKI.

## Discussion

Although Hep G2 (liver cancer – mesodermal tissue origin) and MDA-MB-231 (mammary epithelial adenocarcinoma – ectodermal tissue origin) are different cancer types, they shared a large number of CKI DE genes with similar expression profiles, presumably these shared genes include CKI response genes that are essential to the apoptotic response triggered by CKI. However, the number of shared CKI DE genes is too high to allow straight forward identification of genes critical to the CKI response. We therefore decided to combine these data with previously reported CKI DE genes from MCF-7 cells [28] in order to reduce the number of core CKI response genes. The intersection of MCF-7 CKI DE genes with the shared CKI DE genes yielded 363 core CKI DE genes (Additional file 8: Fig. S4).

Among the 363 core CKI DE genes, cytochrome P450 family 1 subfamily A member 1 (CYP1A1) gene is the most over-expressed. This gene is consistently up-regulated by CKI in all three cell lines, and showed significant dose response. In liver cancer cells, over-expression of CYP1A1 induced by plant natural products has been associated with Aryl-hydrocarbon Receptor transformation [2, 49]. Furthermore, as a steroid-metabolizing enzyme, CYP1A1 is part of cancer metabolic processes relevant to steroid hormone responsive tumors, such as breast cancer, ovarian cancer and prostate cancer [24, 26, 23, 27]. Therefore, CYP1A1 may be of particular interest for understanding the mechanism of action of CKI on cancer cells.

Comparison of the 363 core genes to the 135 Tumor Alterations Relevant for Genomics-driven Therapy (TARGET) genes (version 3) from The Broad Institute (https://www.broadinstitute.org/cancer/cga/target) identified 7 DE genes that were shared across the three cell lines and two time points (Fig. 7a). Of these seven genes, six (TP53, CCND1, MYD88 (Myeloid differentiation primary response gene 88), EWSR1, TMPRSS2 and IDH1 (isocitrate dehydrogenase 1) were similarly regulated (either always over-expressed or under-expressed), while CCND3 was over-expressed in all three cell lines at both time points except at 48 hours in MCF-7 cells, where it was under-expressed.

**Figure 7.**
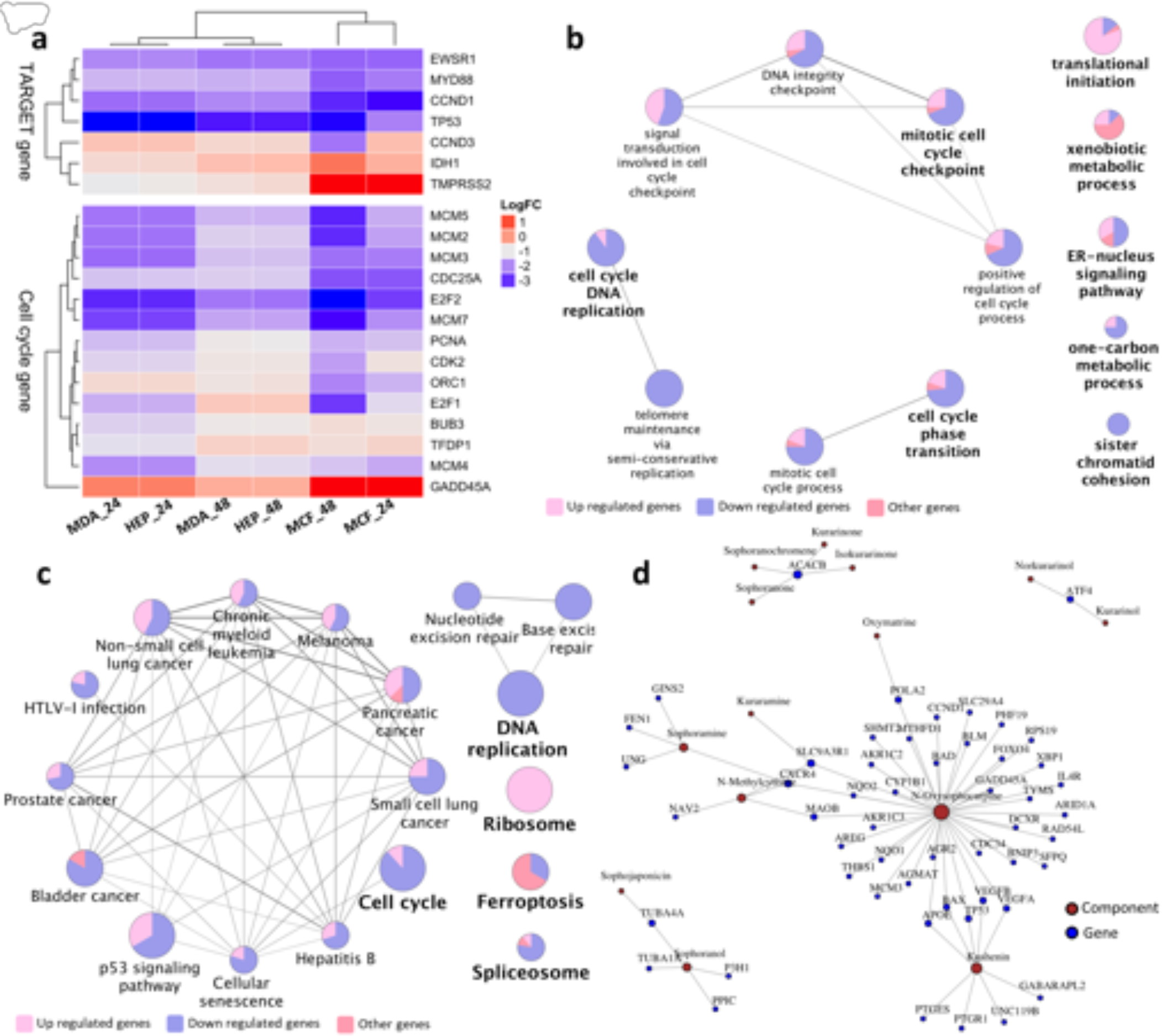
Analysis of CKI regulated core genes from this report combined with previous available data. A) Fold changes of TARGET and cell cycle regulatory gene expression in MDA-MB-231, Hep G2 and MCF-7 [28] cell lines 24 and 48 hours after CKI treatment. Only seven TARGET genes are affected by CKI in all three cell lines. Most of the 14 cell cycle regulatory genes differentially expressed in all three cell lines are down-regulated. B) GO term enrichment analysis of 363 core genes from MDA-MB-231, Hep G2 and MCF-7 cell lines. C) KEGG pathway enrichment of 363 core genes from MDA-MB-231, Hep G2 and MCF-7 cell lines. D) Some individual compounds present in CKI linked to genes they regulate that are also found in this report and our previous study [28]. Node size is proportional to the number of related components/genes.

The TP53 gene encodes a tumor suppressor protein, that can induce apoptosis [11]. However, in all cell lines TP53 was down-regulated, and all cell lines showed increased apoptosis. This suggests that CKI induced apoptosis was not TP53-dependent. Support for this comes from the fact that transcripts for PCNA (proliferating cell nuclear antigen), and a group of transcription factors: MCM (mini-chromosome maintenance) complex and the E2F family are down-regulated. The E2F transcription factors regulate the cell cycle and TP53-dependent and -independent apoptosis [37, 12, 42, 33]. In addition, other core genes present in the TARGET database have also been shown to induce apoptosis. For example, inhibition of MYD88 induces apoptosis in both triple negative breast cancer and bladder cancer [3, 43]. The increased expression of IDH1 may be important, as IDH1 is frequently mutated in cancers [16] and when mutated, it causes loss of *α*-ketoglutarate production and may be important for the Warburg effect. TMPRSS2 (transmembrane protease, serine 2) has also been shown to regulate apoptosis in cancer [1]. Therefore, CKI may induce apoptosis through a variety of means.

In the GO (Fig. 7b) and KEGG (Fig. 7c) over-representation analysis of the 363 core genes yielded enrichment for cell cycle and cancer pathways. In the GO enriched genes, cell cycle and related pathways accounted for the majority of functional sub-clusters. In the KEGG enriched pathways, cell cycle and cancer pathways predominated in a single cluster. Most of the core genes in GO and KEGG clusters were down-regulated by CKI. In addition to the cell cycle, CKI treatment also caused enrichment for terms or pathways related to cancer progression, such as “focal adhesion” and “blood vessel development”. (Additional file 6: Table S4, sheet 5-8). These developmental processes contribute to tumorigenesis and metastasis [20, 36]. It is tempting to speculate that CKI may alter these functions *in vivo*, possibly altering angiogenesis which is critical for tumors [9]. In addition, there were metabolic pathways and terms that were also identified as perturbed by CKI. Effects on many targets/pathways is one of the expected features of TCM drugs which likely hit multiple targets [7].

We have examined the effect of a complex mixture of plant natural products (CKI) on different cancer cell lines and have identified specific, consistent effects on gene expression resulting from this mixture. However, the complexity of CKI makes it difficult to determine the mechanism of action of individual components, and often testing of individual components has resulted in either no effect or contradictory results in the research literature. In spite of this complexity, it is possible to map our results on to a pre-existing corpus of work that links individual natural compounds to changes in gene expression. We have used BATMAN [19], an online TCM database of curated links between compounds and gene expression. Based on this resource, we have identified 14 components of CKI that have been linked to the regulation of 52 of our core genes (Fig. 7d). We can see from the network diagram in Fig. 7d that one to one, one to many and many to many relationships exist between CKI components and genes which is consistent with previous studies [38, 18, 46]. As more information becomes available for individual components, we will be able to construct a more comprehensive model of CKI mechanism based on network analysis.

## Conclusion

Our systematic analysis of gene expression changes in cancer cells caused by a complex herbal extract used in TCM has proven to be effective at identifying candidate molecular pathways. CKI has consistent and specific effects on gene expression across multiple cancer cell lines and it also consistently induces apoptosis *in vitro*. These effects show that CKI can suppress the expression of cell cycle regulatory genes and other well characterized cancer related genes and pathways. Validation of a subset of DE genes at lower doses of CKI has shown a dose-response relationship that suggests that CKI may have similar effects *in vivo* at clinically relevant concentrations. Our results provide a molecular basis for further investigation of the mechanism of action of CKI.

## Authors’ e-mail addresses

Jian Cui

Zhipeng Qu

Yuka Harata-Lee

Hanyuan Shen

Thazin Nwe Aung

Wei Wang

R.Daniel Kortschak

David L Adelson

## Author contributions statement

JC experimental design, carried out experiments, analyzed data, wrote paper, ZQ experimental design, assisted with experiments, assisted with data analysis, wrote paper, YHL experimental design, assisted with experiments, assisted with data analysis, wrote paper, HS assisted with experiments, TNA assisted with experiments, WW assisted with experimental design, assisted with experiments, RDK experimental design, and DLA supervised the research, acquired funding for the experiments, experimental design, wrote paper.

## Data Availability

All RNA-seq data raw and processed data were deposited at the Gene Expression Omnibus (GEO) data repository (XXXXXXX).

## Abbreviations

DE: differentially expressed
TCM: traditional Chinese medicine
CKI: compound Kushen injection
GO: Gene Ontology
DO: Disease Ontology
KEGG: Kyoto Encyclopedia of Gene and Genomes
PI: propidium iodide.

## References

[1] Afar, D. E., Vivanco, I., Hubert, R. S., Kuo, J., Chen, E., Saffran, D. C., Raitano, A. B., Jakobovits, A., 2001. Catalytic cleavage of the androgen-regulated tmprss2 protease results in its secretion by prostate and prostate cancer epithelia. Cancer research 61 (4), 1686–1692.

[2] Anwar-Mohamed, A., El-Kadi, A. O., 2009. Sulforaphane induces cyp1a1 mrna, protein, and catalytic activity levels via an ahr-dependent pathway in murine hepatoma hepa 1c1c7 and human hepg2 cells. Cancer letters 275 (1), 93–101.

[3] Christensen, A. G., Ehmsen, S., Terp, M. G., Batra, R., Alcaraz, N., Baumbach, J., Noer, J. B., Moreira, J., Leth-Larsen, R., Larsen, M. R., 2017. Elucidation of altered pathways in tumor-initiating cells of triple-negative breast cancer: A useful cell model system for drug screening. STEM CELLS.

[4] Csardi, G., Nepusz, T., 2006. The igraph software package for complex network research. InterJournal, Complex Systems 1695 (5), 1–9.

[5] Cui, M., Li, H., Hu, X., 2014. Similarities between “big data” and traditional chinese medicine information. Journal of Traditional Chinese Medicine 34 (4), 518–522.

[6] Dobin, A., Davis, C. A., Schlesinger, F., Drenkow, J., Zaleski, C., Jha, S., Batut, P., Chaisson, M., Gingeras, T. R., 2013. Star: ultrafast universal rna-seq aligner. Bioinformatics 29 (1), 15–21.

[7] Efferth, T., Li, P. C., Konkimalla, V. S. B., Kaina, B., 2007. From traditional chinese medicine to rational cancer therapy. Trends in molecular medicine 13 (8), 353–361.

[8] Gao, L., Wang, K.-x., Zhou, Y.-z., Fang, J.-s., Qin, X.-m., Du, G.-h., 2018. Uncovering the anticancer mechanism of compound kushen injection against hcc by integrating quantitative analysis, network analysis and experimental validation. Scientific reports 8 (1), 624.

[9] Hanahan, D., Folkman, J., 1996. Patterns and emerging mechanisms of the angiogenic switch during tumorigenesis. cell 86 (3), 353–364.

[10] Hao, D. C., Xiao, P. G., 2014. Network pharmacology: a rosetta stone for traditional chinese medicine. Drug development research 75 (5), 299–312.

[11] Harris, S. L., Levine, A. J., 2005. The p53 pathway: positive and negative feedback loops. Oncogene 24 (17), 2899.

[12] Hollern, D. P., Honeysett, J., Cardiff, R. D., Andrechek, E. R., 2014. The e2f transcription factors regulate tumor development and metastasis in a mouse model of metastatic breast cancer. Molecular and cellular biology 34 (17), 3229–3243.

[13] Jiang, W.-Y., 2005. Therapeutic wisdom in traditional chinese medicine: a perspective from modern science. Trends in pharmacological sciences 26 (11), 558–563.

[14] Kamburov, A., Wierling, C., Lehrach, H., Herwig, R., 2008. Consensuspathdb—a database for integrating human functional interaction networks. Nucleic acids research 37 (suppl 1), D623–D628.

[15] Kim, D., Pertea, G., Trapnell, C., Pimentel, H., Kelley, R., Salzberg, S. L., 2013. Tophat2: accurate alignment of transcriptomes in the presence of insertions, deletions and gene fusions. Genome Biol 14 (4), R36.

[16] Li, X., Xu, X., Wang, J., Yu, H., Wang, X., Yang, H., Xu, H., Tang, S., Li, Y., Yang, L., 2012. A system-level investigation into the mechanisms of chinese traditional medicine: Compound danshen formula for cardiovascular disease treatment. PloS one 7 (9), e43918.

[17] Liao, C.-Y., Lee, C.-C., Tsai, C.-c., Hsueh, C.-W., Wang, C.-C., Chen, I., Tsai, M.-K., Liu, M.-Y., Hsieh, A.-T., Su, K.-J., 2015. Novel investigations of flavonoids as chemopreventive agents for hepatocellular carcinoma. BioMed research international 2015.

[18] Liu, Y., Xu, Y., Ji, W., Li, X., Sun, B., Gao, Q., Su, C., 2014. Anti-tumor activities of matrine and oxymatrine: literature review. Tumor Biology 35 (6), 5111–5119.

[19] Liu, Z., Guo, F., Wang, Y., Li, C., Zhang, X., Li, H., Diao, L., Gu, J., Wang, W., Li, D., 2016. Batman-tcm: a bioinformatics analysis tool for molecular mechanism of traditional chinese medicine. Scientific reports 6.

[20] Luo, M., Guan, J.-L., 2010. Focal adhesion kinase: a prominent determinant in breast cancer initiation, progression and metastasis. Cancer letters 289 (2), 127–139.

[21] Luo, W., Brouwer, C., 2013. Pathview: an r/bioconductor package for pathway-based data integration and visualization. Bioinformatics 29 (14), 1830–1831.

[22] Ma, X., Li, R.-S., Wang, J., Huang, Y.-Q., Li, P.-Y., Wang, J., Su, H.-B., Wang, R.-L., Zhang, Y.-M., Liu, H.-H., 2016. The therapeutic efficacy and safety of compound kushen injection combined with transarterial chemoembolization in unresectable hepatocellular carcinoma: an update systematic review and meta-analysis. Frontiers in pharmacology 7.

[23] Mitsui, Y., Chang, I., Kato, T., Hashimoto, Y., Yamamura, S., Fukuhara, S., Wong, D. K., Shiina, M., Imai-Sumida, M., Majid, S., 2016. Functional role and tobacco smoking effects on methylation of cyp1a1 gene in prostate cancer. Oncotarget 7 (31), 49107.

[24] Nandekar, P. P., Khomane, K., Chaudhary, V., Rathod, V. P., Borkar, R. M., Bhandi, M. M., Srinivas, R., Sangamwar, A. T., Guchhait, S. K., Bansal, A. K., 2016. Identification of leads for antiproliferative activity on mda-mb-435 human breast cancer cells through pharmacophore and cyp1a1-mediated metabolism. European journal of medicinal chemistry 115, 82–93.

[25] Ninomiya, M., Koketsu, M., 2013. Minor flavonoids (chalcones, flavanones, dihydrochalcones, and aurones). Springer, pp. 1867–1900.

[26] Ou, C., Zhao, Y., Liu, J.-H., Zhu, B., Li, P.-Z., Zhao, H.-L., 2016. Relationship between aldosterone synthase cyp1a1 mspi gene polymorphism and prostate cancer risk. Technology in cancer research & treatment, 1533034615625519.

[27] Piotrowska-Kempisty, H., Klupczyńska, A., Trzybulska, D., Kulcenty, K., Sulej-Suchomska, A. M., Kucińska, M., Mikstacka, R., Wierzchowski, M., Murias, M., Baer-Dubowska, W., 2017. Role of cyp1a1 in the biological activity of methylated resveratrol analogue, 3, 4, 5, 4’-tetramethoxystilbene (dmu-212) in ovarian cancer a-2780 and non-cancerous hose cells. Toxicology letters 267, 59–66.

[28] Qu, Z., Cui, J., Harata-Lee, Y., Aung, T. N., Feng, Q., Raison, J. M., Kortschak, R. D., Adelson, D. L., 2016. Identification of candidate anti-cancer molecular mechanisms of compound kushen injection using functional genomics. Oncotarget 7 (40), 66003–66019.

[29] Robinson, M. D., McCarthy, D. J., Smyth, G. K., 2010. edger: a bioconductor package for differential expression analysis of digital gene expression data. Bioinformatics 26 (1), 139–140.

[30] Schriml, L. M., Arze, C., Nadendla, S., Chang, Y.-W. W., Mazaitis, M., Felix, V., Feng, G., Kibbe, W. A., 2011. Disease ontology: a backbone for disease semantic integration. Nucleic acids research 40 (D1), D940–D946.

[31] Shu, G., Yang, J., Zhao, W., Xu, C., Hong, Z., Mei, Z., Yang, X., 2014. Kurarinol induces hepatocellular carcinoma cell apoptosis through suppressing cellular signal transducer and activator of transcription 3 signaling. Toxicology and applied pharmacology 281 (2), 157–165.

[32] Song, Y.-N., Zhang, G.-B., Zhang, Y.-Y., Su, S.-B., 2013. Clinical applications of omics technologies on zheng differentiation research in traditional chinese medicine. Evidence-Based Complementary and Alternative Medicine 2013.

[33] Sun, H., Xu, Y., Yang, X., Wang, W., Bai, H., Shi, R., Nayar, S. K., Devbhandari, R. P., He, Y., Zhu, Q., 2013. Hypoxia inducible factor 2 alpha inhibits hepatocellular carcinoma growth through the transcription factor dimerization partner 3/e2f transcription factor 1–dependent apoptotic pathway. Hepatology 57 (3), 1088–1097.

[34] Wang, P., Chen, Z., 2013. Traditional chinese medicine zheng and omics convergence: A systems approach to post-genomics medicine in a global world. Omics: a journal of integrative biology 17 (9), 451–459.

[35] Wang, W., You, R.-l., Qin, W.-j., Hai, L.-n., Fang, M.-j., Huang, G.-h., Kang, R.-x., Li, M.-h., Qiao, Y.-f., Li, J.-w., 2015. Anti-tumor activities of active ingredients in compound kushen injection. Acta Pharmacologica Sinica 36 (6), 676–679.

[36] Warren, R. S., Yuan, H., Matli, M. R., Gillett, N. A., Ferrara, N., 1995. Regulation by vascular endothelial growth factor of human colon cancer tumorigenesis in a mouse model of experimental liver metastasis. The Journal of clinical investigation 95 (4), 1789–1797.

[37] Woods, K., Thomson, J. M., Hammond, S. M., 2007. Direct regulation of an oncogenic micro-rna cluster by e2f transcription factors. Journal of Biological Chemistry 282 (4), 2130–2134.

[38] Wu, C., Huang, W., Guo, Y., Xia, P., Sun, X., Pan, X., Hu, W., 2015. Oxymatrine inhibits the proliferation of prostate cancer cells in vitro and in vivo. Molecular medicine reports 11 (6), 4129–4134.

[39] Yang, X., Huang, M., Cai, J., Lv, D., Lv, J., Zheng, S., Ma, X., Zhao, P., Wang, Q., 2017. Chemical profiling of anti-hepatocellular carcinoma constituents from caragana tangutica maxim. by off-line semi-preparative hplc-nmr. Natural product research 31 (10), 1150–1155.

[40] Yu, G., Wang, L.-G., Han, Y., He, Q.-Y., 2012. clusterprofiler: an r package for comparing biological themes among gene clusters. OMICS: a Journal of Integrative Biology 16 (5), 284–287. URL http://www.ncbi.nlm.nih.gov/pmc/articles/PMC3339379/

[41] Yu, P., Liu, Q., Liu, K., Yagasaki, K., Wu, E., Zhang, G., 2009. Matrine suppresses breast cancer cell proliferation and invasion via vegf-akt-nf-*κ*b signaling. Cytotechnology 59 (3), 219–229.

[42] Zaldua, N., Llavero, F., Artaso, A., Gálvez, P., Lacerda, H. M., Parada, L. A., Zugaza, J. L., 2016. Rac1/p21-activated kinase pathway controls retinoblastoma protein phosphorylation and e2f transcription factor activation in b lymphocytes. FEBS journal 283 (4), 647–661.

[43] Zhang, H., Ye, Y., Li, M., Ye, S., Huang, W., Cai, T., He, J., Peng, J., Duan, T., Cui, J., 2016. Cxcl2/mif-cxcr2 signaling promotes the recruitment of myeloid-derived suppressor cells and is correlated with prognosis in bladder cancer. Oncogene.

[44] Zhang, J.-Q., Li, Y.-M., Liu, T., He, W.-T., Chen, Y.-T., Chen, X.-H., Li, X., Zhou, W.-C., Yi, J.-F., Ren, Z.-J., 2010. Antitumor effect of matrine in human hepatoma g2 cells by inducing apoptosis and autophagy. World journal of gastroenterology: WJG 16 (34), 4281.

[45] Zhang, L. P., Jiang, J. K., Tam, J. W. O., Zhang, Y., Liu, X. S., Xu, X. R., Liu, B. Z., He, Y. J., 2001. Effects of matrine on proliferation and differentiation in k-562 cells. Leukemia Research 25 (9), 793–800. URL http://www.sciencedirect.com/science/article/pii/S0145212600001454

[46] Zhang, X., Yu, H., 2016. Matrine inhibits diethylnitrosamine-induced hcc proliferation in rats through inducing apoptosis via p53, bax-dependent caspase-3 activation pathway and down-regulating mlck overexpression. Iranian Journal of Pharmaceutical Research: IJPR 15 (2), 491.

[47] Zhang, X.-L., Cao, M.-A., Pu, L.-P., Huang, S.-S., Gao, Q.-X., Yuan, C.-S., Wang, C.-M., 2013. A novel flavonoid isolated from sophora flavescens exhibited anti-angiogenesis activity, decreased vegf expression and caused g0/g1 cell cycle arrest in vitro. Die Pharmazie-An International Journal of Pharmaceutical Sciences 68 (5), 369–375.

[48] Zhao, Z., Fan, H., Higgins, T., Qi, J., Haines, D., Trivett, A., Oppenheim, J. J., Wei, H., Li, J., Lin, H., 2014. Fufang kushen injection inhibits sarcoma growth and tumor-induced hyperalgesia via trpv1 signaling pathways. Cancer letters 355 (2), 232–241.

[49] Zhou, Y., Li, Y., Zhou, T., Zheng, J., Li, S., Li, H.-B., 2016. Dietary natural products for prevention and treatment of liver cancer. Nutrients 8 (3), 156.

