## Supplementary material for "The Effect of Compound Kushen Injection on Cancer Cells: Integrated Identification of Candidate Molecular Mechanisms": Supplementary_table_figure_legends.pdf

**Additional file 1: Table S1. RT-qPCR target genes and their primer sequences.**

**Additional file 2: Figure S1. MDS plot of the DE gene distribution of two cell lines under different conditions.**

**Additional file 3: Table S2. Mapping rate of each cell line.**

**Additional file 4: Table S3. List of DE genes in each cell line at each time point.**

**Additional file 5: Figure S2. The GO semantic similarity analysis of each data set.** (A-D) Each small square represents a GO biological process functions in level 3. Size of the squares positively correlates with the statistical significance of related biological process. Different colour is to distinguish biological process clusters that are descripted by the top shaded functional representatives.

**Additional file 6: Table S4. The summary of functional analysis of both separate datasets and shared datasets.**

- **Sheet 1-4: GO enrichment of each cell line at two time points.** Selection standard: cut off $p$ value<0.01, cut off $q$ value<0.01.

- **Sheet 5-8: KEGG enrichment of each cell line at two time points.** Selection standard: cut off $p$ value<0.01, cut off $q$ value<0.01.

- **Sheet 10-12: DO enrichment of each cell line at two time points.** Selection standard: cut off $p$ value<0.01, cut off $q$ value<0.01.

- **Sheet 13: GO enrichment of shared genes by both cell lines.** Selection standard: cut off $p$ value<0.01.

- **Sheet 14: KEGG enrichment of shared genes by both cell lines.** Selection standard: cut off $p$ value<0.01.

**Additional file 7: Figure S3. DE genes distribution of two cell lines in the pathways in cancer.** In the cell cycle pathway, each coloured box is separated into 4 parts, from left to right representing 24h CKI treated Hep G2, 48h CKI treated Hep G2, 24h CKI treated MDA-MB-231 and 48h CKI treated MDA-MB-231.

**Additional file 8: Figure S4. The heatmap of core genes of three cell lines.** Heatmap revealing the expression fold changes of core genes in three cell lines at two time points. All the core genes can be separated into 3 clusters, namely consistently up/down regulated genes and uneven genes.
