## Supplementary figures and images for "The Effect of Compound Kushen Injection on Cancer Cells: Integrated Identification of Candidate Molecular Mechanisms"

### Additional file 2_figure S1.tif

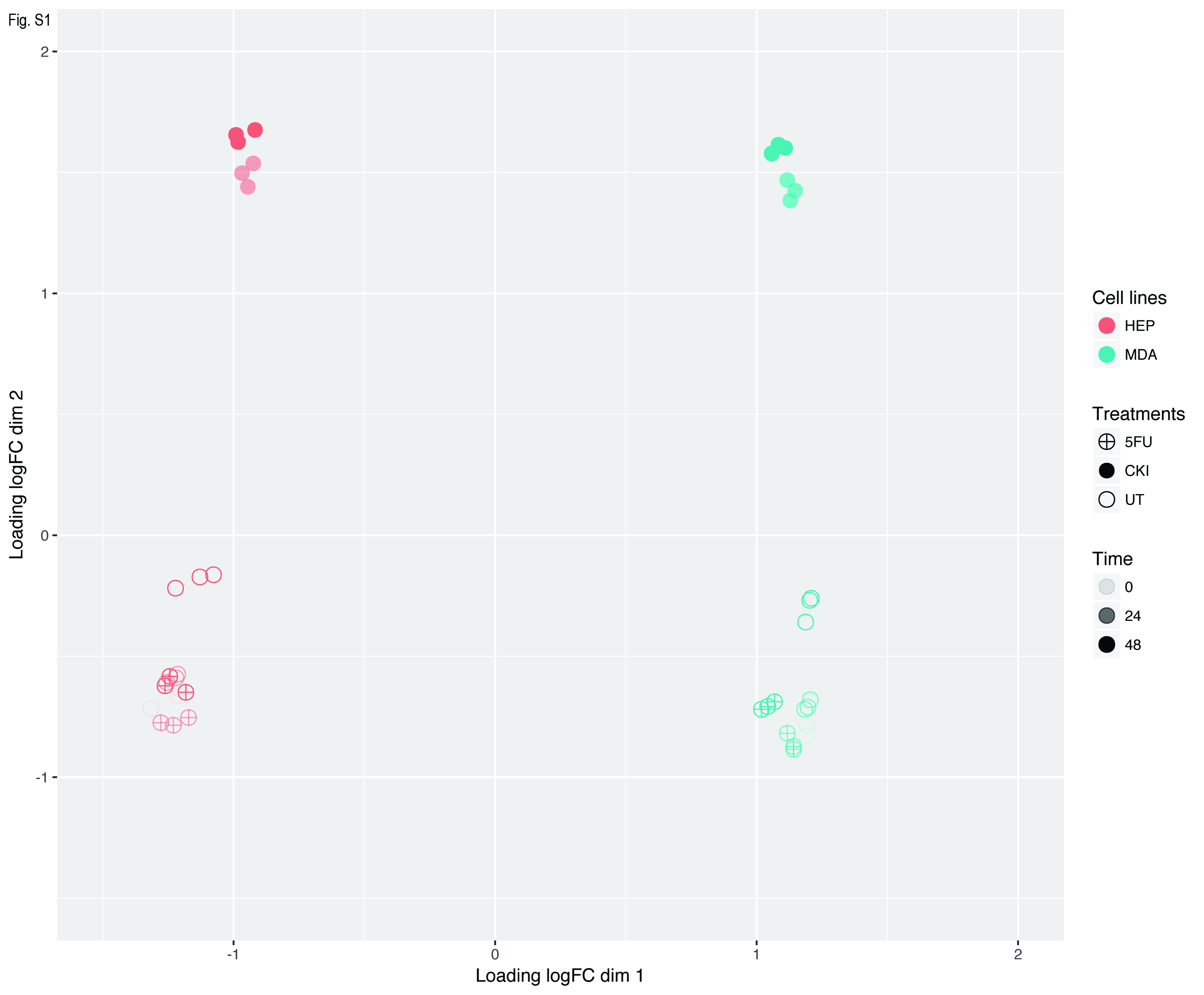

### Additional file 5_figure S2.pdf

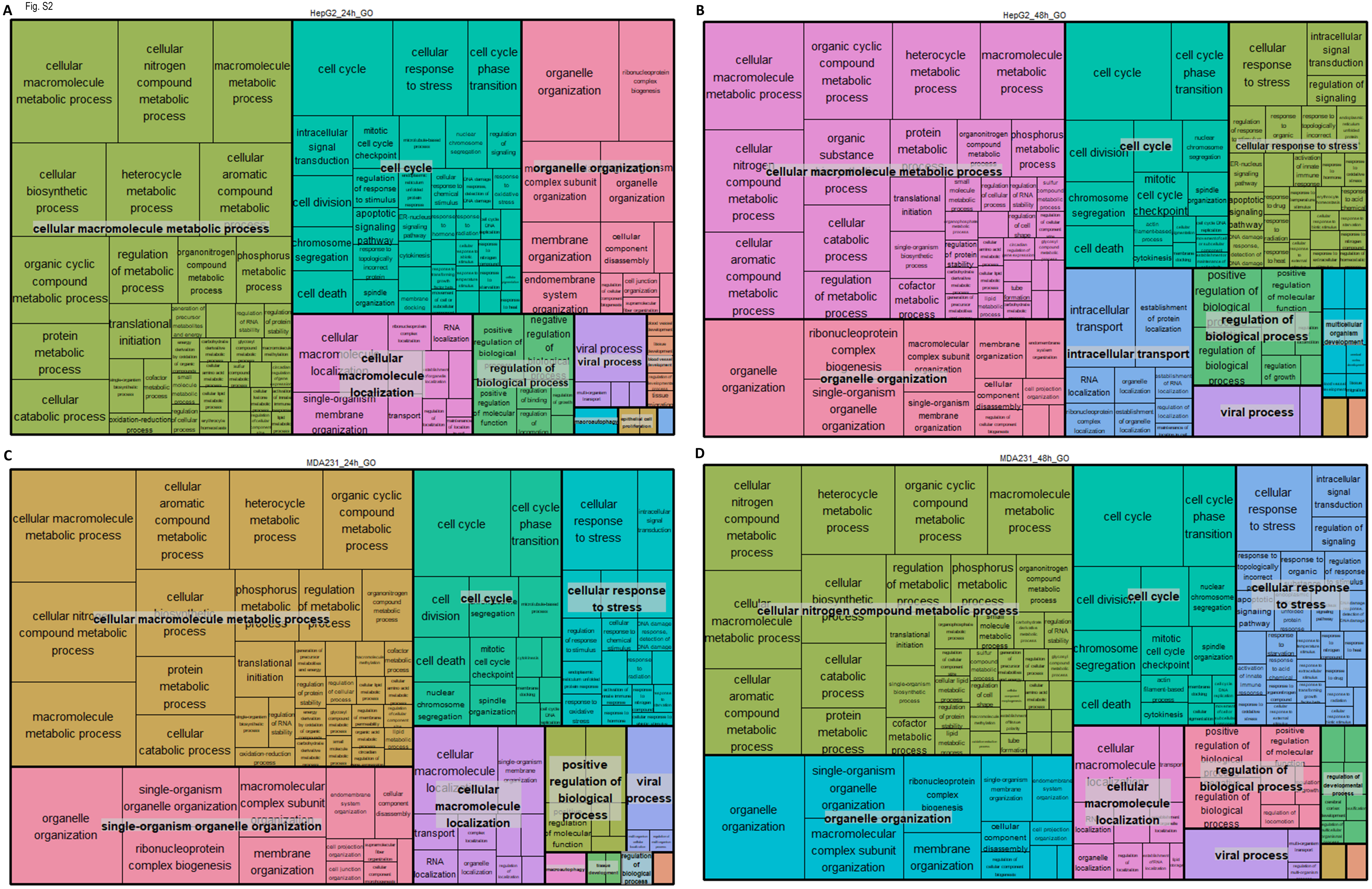

### Additional file 7_figure S3.tif

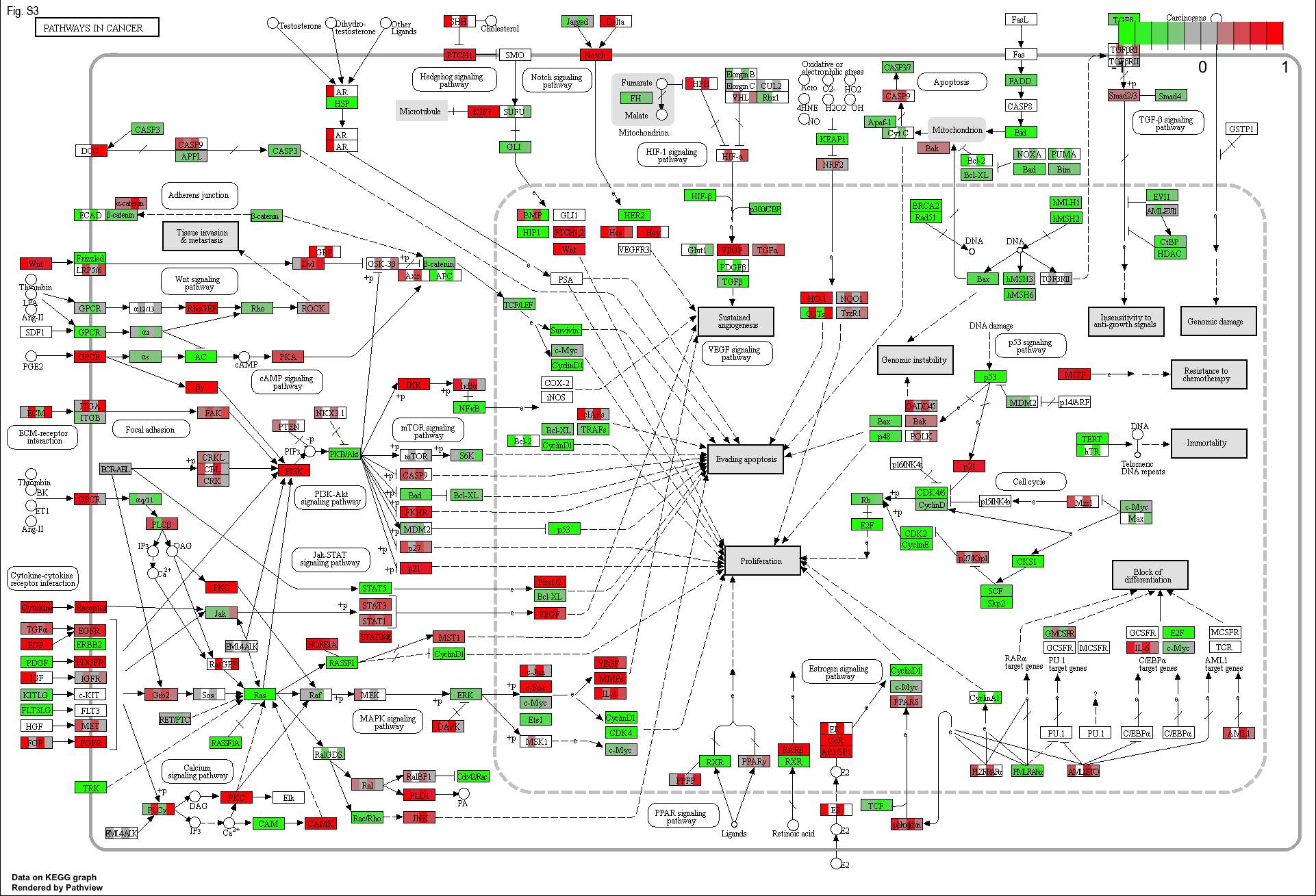

### Additional file 8_figure S4.tif

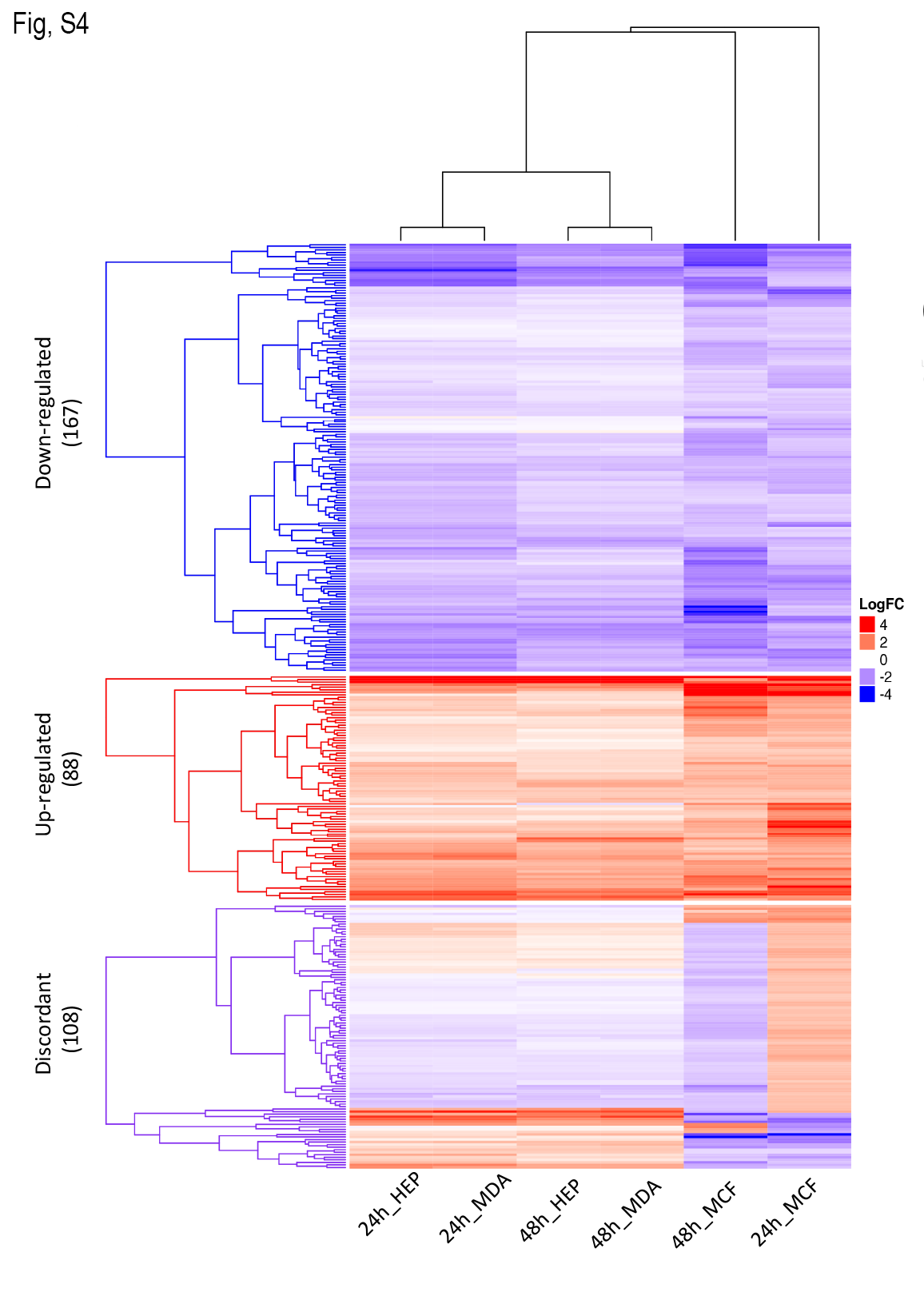
